# Variability of bacterial behavior in the mammalian gut captured using a growth-linked single-cell synthetic gene oscillator

**DOI:** 10.1101/472720

**Authors:** David T Riglar, David L Richmond, Laurent Potvin-Trottier, Andrew A Verdegaal, Alexander D Naydich, Somenath Bakshi, Emanuele Leoncini, Johan Paulsson, Pamela A Silver

**Author notes:** Authors contributed equally (David L Richmond, Laurent Potvin-Trottier).

## Abstract

The dynamics of the bacterial population that comprises the gut microbiota plays key roles in overall mammalian health. However, a detailed understanding of bacterial growth within the gut is limited by the inherent complexity and inaccessibility of the gut environment. Here, we deploy an improved synthetic genetic oscillator to investigate dynamics of bacterial colonization and growth in the mammalian gut under both healthy and disease conditions. The synthetic oscillator, when introduced into both *Escherichia coli* and *Salmonella* Typhimurium maintains regular oscillations with a constant period in generations across growth conditions. We determine the phase of oscillation from individual bacteria using image analysis of resultant colonies and thereby infer the number of cell divisions elapsed. In doing so, we demonstrate robust functionality and controllability of the oscillator circuit’s activity during bacterial growth *in vitro*, in a simulated murine gut microfluidic environment, and *in vivo* within the mouse gut. We determine different dynamics of bacterial colonization and growth in the gut under normal and inflammatory conditions. Our results show that a precise genetic oscillator can function in a complex environment and reveal single cell behavior under diverse conditions where disease may create otherwise impossible-to-quantify variability in growth across the population.

## Introduction

The mammalian gut microbiota’s composition and function are critical for immune development, nutrition, and the maintenance of health. Indeed, dysbiosis – an altered state of the microbiota indicative of perturbations to bacterial growth rates, death rates, or the carrying capacity of the environment– has been linked to an increasing array of diseases^1^

Direct measurement of bacterial growth under both normal and disease conditions is an ongoing challenge. Due to the heterogeneity of niches within the mammalian gut, a comprehensive understanding of growth requires single-cell measurements and an ability to distinguish division independently from death and elimination. The dynamics of change within the microbiota are also rapid; some alterations are measurable hours after dietary change and infection^2,3^.

Oscillators are extensively utilized as clocks and timers in computation and biology, including control of the cell cycle and circadian clocks^4,5^, and are thus a good option for growth-related applications. Synthetic gene oscillators in living cells have been refined over past decades^6–15^, led by the development of the repressilator circuit – a simple negative feedback loop constructed from three transcriptional repressor proteins which inhibit each other in turn (Tn10 TetR, bacteriophage λ CI and *E. coli* LacI)^6^.

The original repressilator circuit was recently redesigned using stochastic modelling to afford a highly regular and robust oscillator with reduced error propagation and information losses, referred to herein as the repressilator 2.0 (Fig 1a; Supplementary Fig 1a)^8^. The incorporation of extra repressor binding sites on a ‘sponge plasmid’ was a key element in reducing oscillation variability, with the circuit keeping phase in single cells for hundreds of bacterial generations^8^. The repressilator 2.0 period is growth-rate independent and linked to bacterial divisions^8^. Due to the circuit’s low variance between cells, and its independence from external feedback and entrainment cues, it represents an attractive timer for use in complex environments.

**Figure 1:**
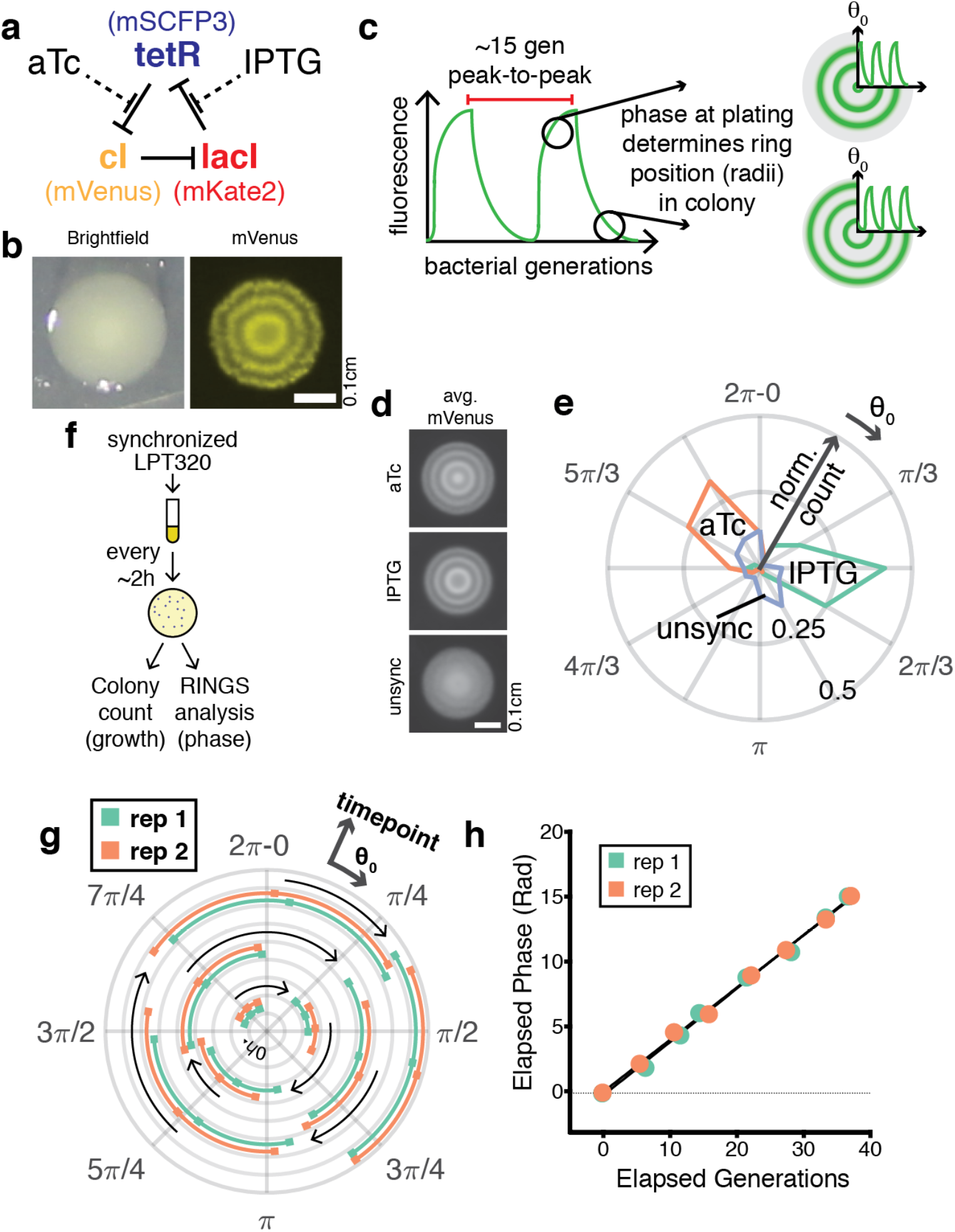
RINGS analysis measures phase of colony-initiating bacteria carrying the repressilator 2.0. **a.** The repressilator comprises 3 repressors, that act on expression of each other (and a corresponding fluorescent reporter) in turn, leading to fluorescent oscillations. **b.** Colonies expressing the repressilator 2.0 display fluorescent rings controlled by the circuit’s oscillations. **c.** Repressilator phase progresses based on bacterial divisions, with an ~15 generation period. The phase of the colony-initiating bacterium (Θ_0_) controls the position of the fluorescent rings forming within that colony. **d**. LPT239 bacteria, carrying the repressilator, are synchronized by exposure to aTc or IPTG compared to unsynchronized controls as demonstrated by average projections of aligned colony images for the population (IPTG n=93; aTc n=62; unsync n= 49). Scale = 0.1cm. **e**. RINGS analysis demonstrates the ability of the repressilator to report on bacterial phase. Graph shows polar histogram representing (radius) normalized counts of colonies within a (angle) given Θ_0_ range. **f**. LPT320 bacteria were synchronized with aTc and grown in log-phase growth at 37°C, plating samples for colony counts and RINGS analysis ~2 hourly. **g**. RINGS analysis demonstrated that population phase (Θ_0_) shifted throughout the experiment. Graph shows circular mean ± angular deviation for two replicates, with growth corresponding to a clockwise phase shift. Each ring corresponds to a discrete timepoint. **h**. Repressilator phase correlates with the estimated generations as determined by CFU counts from agar plating. Graph shows mean with 95% CI (error bars smaller than datapoints) of elapsed phase after removal of aTc.

Here, we use the repressilator 2.0 to quantify growth on a single-cell level. We infer the phase of a bacterium through image analysis of reporter-gene oscillations within the bacterial colony that it seeds. In this way, we investigate bacterial population dynamics during colonization of, and stable growth within, the mammalian gut. Our results show increased growth variability within the gut under inflamed conditions, demonstrating the importance of single-cell methods for reliable bacterial growth analysis. We also reveal robust functionality and controllability of the repressilator 2.0 across diverse host and environmental contexts, demonstrating its power to drive growth-linked applications in real-world settings.

## Results

### Repressilator function across diverse conditions and bacterial species

Because bacterial colonies expand radially in a uniform manner, with division only occurring at the periphery^16^, synchronous repressilator 2.0 oscillations create stable macroscopic fluorescent rings expressed by bacteria arrested in growth at the colony center (Fig 1b)^8^. The radius of the rings is determined by the phase of the bacterium that seeded the colony (Fig 1c). Using this behavior, we developed a workflow for bacterial colony image capture and processing, which we call <u>R</u>epressilator-based <u>IN</u>ference of <u>G</u>rowth at <u>S</u>ingle-cell level (RINGS), to estimate the phase of individual bacteria at the time of plating (Θ_0_), by fitting the rings within each colony (see Methods; Supplementary Fig 1b).

To demonstrate the utility of the RINGS workflow, *E.coli* LPT239 repressilator 2.0 expressing bacteria (Supplementary Table 1) were phase-synchronized by growth in the presence of Isopropyl β-D-1-thiogalactopyranoside (IPTG) or anhydrotetracycline (aTc), which interrupt repression by LacI and TetR respectively (Fig 1a). Analysis of YFP fluorescence by colony-level imaging (Fig 1d) and RINGS analysis (Fig 1e) demonstrated the ability for RINGS to successfully distinguish between distinct oscillator phases. RINGS analysis was then further optimized by provision of sponge-plasmid binding sites for each repressor (LPT320), which resulted in more consistent fluorescent rings within colonies and allowed analysis using combined CFP and YFP fluorescent data (see Methods; Supplementary Fig 2).

RINGS analysis infers bacterial growth through phase measurement. We grew aTc-synchronized *E. coli* LPT320 bacteria in culture with back dilution to produce constant log-phase growth. The population was sampled by plating at ~2-hour intervals, which were imaged and analyzed using RINGS to measure repressilator phase, and for colony forming unit (CFU) counts to measure average bacterial growth (Fig 1f). Repressilator phase progressed (Fig 1g) according to bacterial growth of the population (Fig 1h), validating the RINGS approach. The gradual increase in the population’s angular deviation also indicated the method’s suitability for longer-term growth measurement (Supplementary Fig 2c)

To expand the potential utility of the repressilator, we tested the circuit’s ability to oscillate across a range of *E. coli* strains (MG1655 – PAS715; and the human probiotic strain, Nissle 1917 – PAS717) and in *S.* Typhimurium LT2 – PAS716 (Supplementary Table 1). Imaging identified fluorescent rings within colonies (Fig 2a-c). RINGS analysis of synchronized PAS715 and PAS716 bacteria over 6 hours of log-phase growth also tracked bacterial divisions (CFU counts), further demonstrating the power of RINGS across these strains (Fig 2d-e).

**Figure 2:**
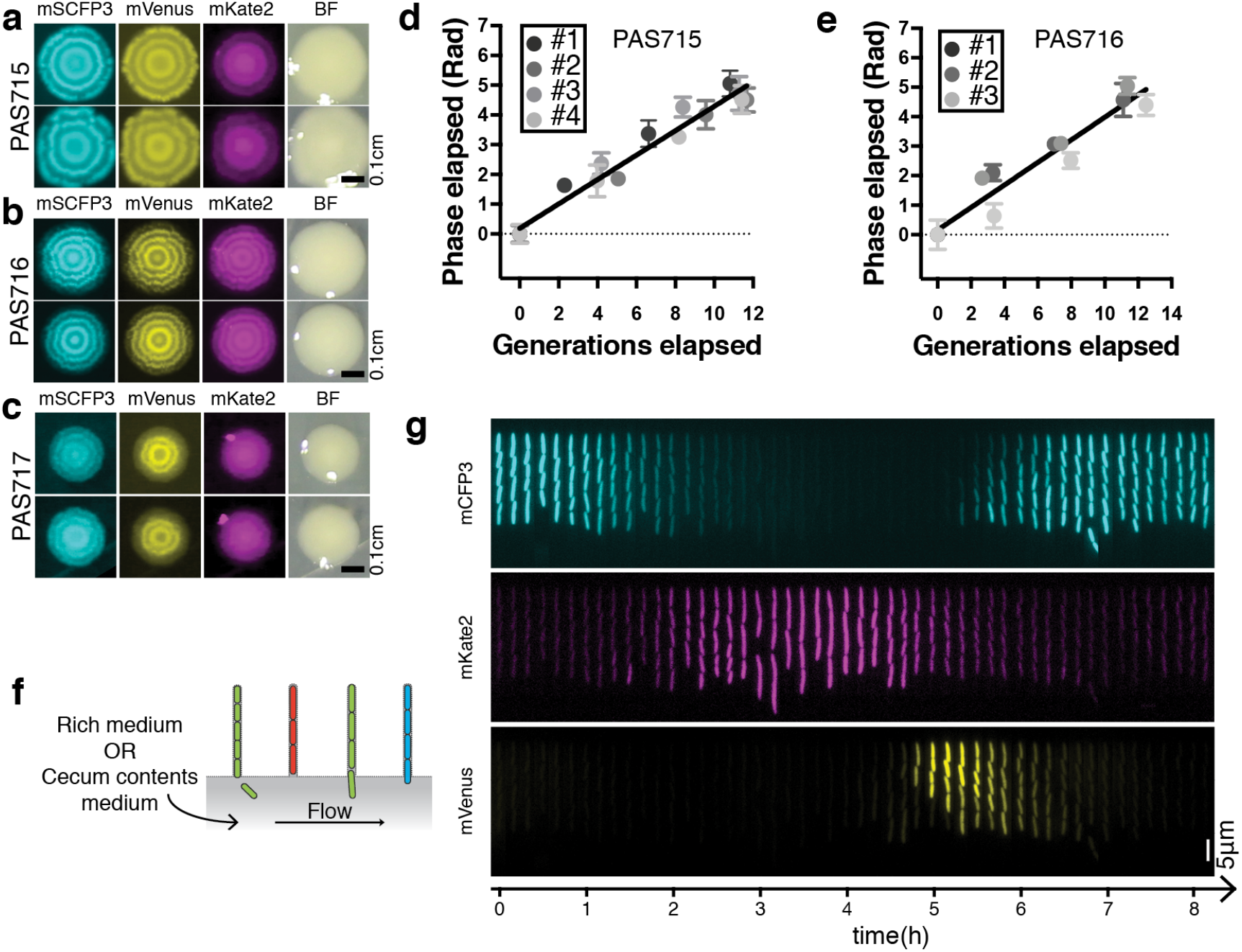
The repressilator 2.0 functions across diverse conditions and bacterial species. **a-c.** Fluorescence and white light images of 2 representative colonies of PAS715 (*E. coli* MG1655), PAS716 (*S*. Typhimurium LT2) and PAS717 (*E. coli* Nissle 1917). Scale bars = 0.1cm. **d-e.** When grown in log-phase liquid culture, repressilator phase progression correlated with the estimated generations elapsed as determined by CFU counts. Graph shows mean with 95% CI of elapsed phase after IPTG removal. **f.** The mother machine is a microfluidic device consisting of trenches that can be seeded with individual bacterial lineages, arranged around a central flow channel that delivers growth medium to the cells. **g.** Kymograph of fluorescent timelapse images from a single growth trench demonstrates oscillation of the repressilator 2.0 in PAS715 bacteria during growth on mouse cecum contents medium. Scale bar = 5μm.

Repressilator 2.0 oscillation periods are robust to both strain and environmental variations. The period of the repressilator was determined either using the regression of RINGS vs growth curves (Fig 2d-e) or during growth in a mother machine – a microfluidic device capable of trapping cells for fluorescent microscopic analysis (Fig 2f) over multiple repressilator cycles (Fig 2g)^17^ Both methods quantified the same period for *E. coli* PAS715 bacteria grown in rich medium (RINGS: 15.3 ± 0.3 SEM gen/period; mother machine: 15.3 ± 0.2 SEM gen/period) (Fig 2d; Table 1), with a similar period calculated for *S.* Typhimurium PAS716 bacteria in the mother machine (16.4 ± 0.6 SEM gen/period) (Fig 2e). PAS715 period length remained unperturbed during growth on extracted mouse cecum contents, used to simulate aspects of the gut environment (Table 1). Similarly, RINGS analysis was able to track variable growth conditions, such as perturbations caused by the antibiotic novobiocin (Supplementary Fig 3). Together, these results indicate that the mechanisms for repressilator 2.0 oscillation are insensitive to the gene regulation, cell-size and stress variations expected between these strains and environmental conditions.

**Table 1:**
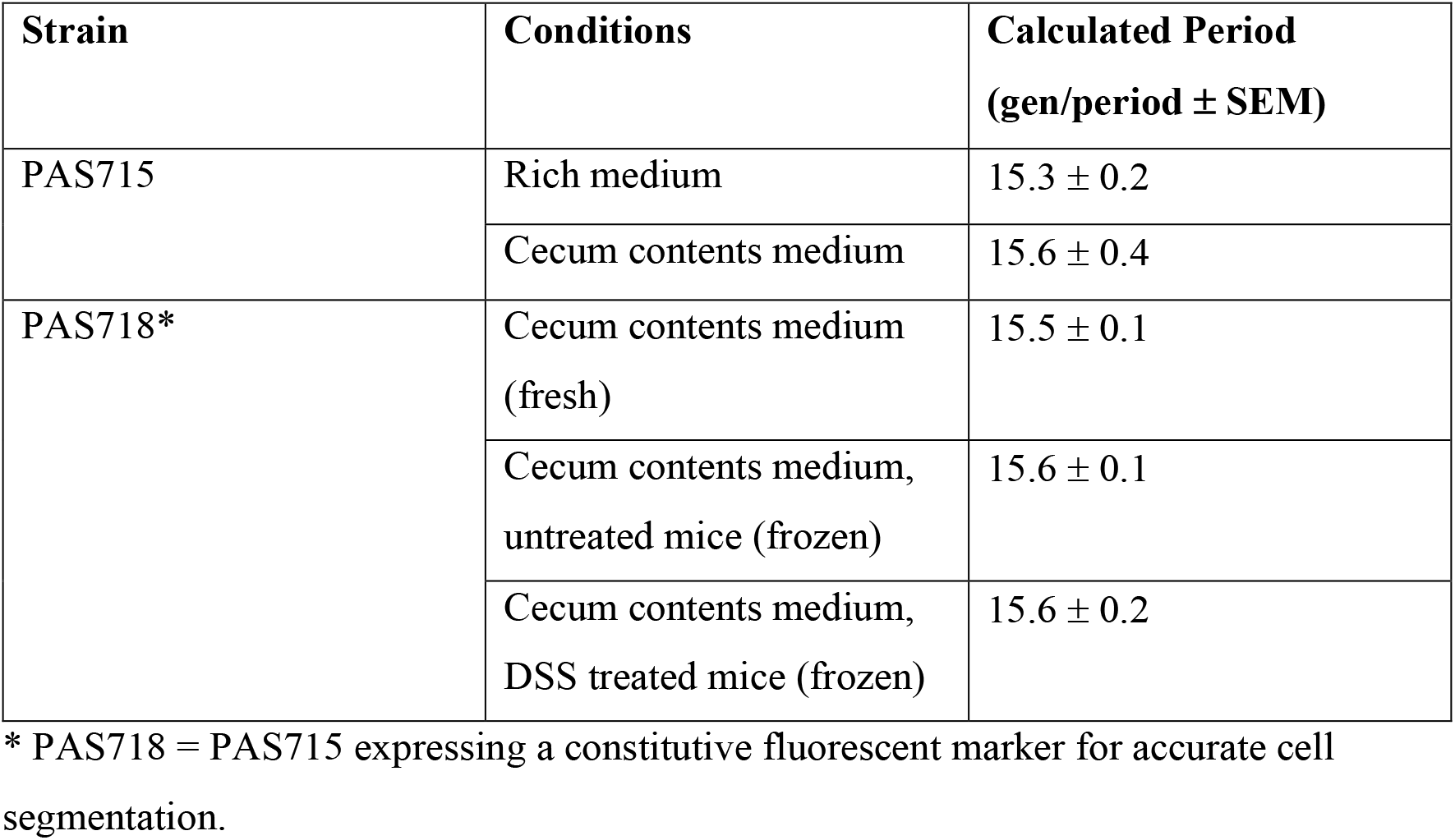
Repressilator period lengths calculated by growth in the mother machine.

| Strain | Conditions | Calculated Period<br>(gen/period ± SEM) |
| --- | --- | --- |
| PAS715 | Rich medium | $15.3 \pm 0.2$ |
| | Cecum contents medium | $15.6 \pm 0.4$ |
| PAS718* | Cecum contents medium<br>(fresh) | $15.5 \pm 0.1$ |
| | Cecum contents medium,<br>untreated mice (frozen) | $15.6 \pm 0.1$ |
| | Cecum contents medium,<br>DSS treated mice (frozen) | $15.6 \pm 0.2$ |

\* PAS718 = PAS715 expressing a constitutive fluorescent marker for accurate cell segmentation.

### Tracking bacterial growth in the mouse gut

To determine the utility of RINGS analysis for the estimate of both growth and population dynamics within the mouse gut, we determined growth of *E. coli* bacteria in the presence or absence of colonization resistance using the streptomycin-treated mouse model. Mice (n=3 per group) were treated with either streptomycin (5 mg/mouse) or saline by oral gavage. 24 hours later, IPTG-synchronized PAS715 were provided to the mice by oral gavage (Fig 3a). Growth of fecal samples on selective plates and RINGS analysis followed phase progression of the population during early colonization of the mouse gut. The phase of PAS715 in both streptomycin treated and untreated mice remained coherent for the entirety of the ~24 hour experiment, indicating similar total growth across each population (Fig 3b-c). Phase progression was greater in streptomycin treated mice than untreated controls, mirroring the higher bacterial loads in these samples (4-8 orders of magnitude difference) (Fig 3d).

**Figure 3:**
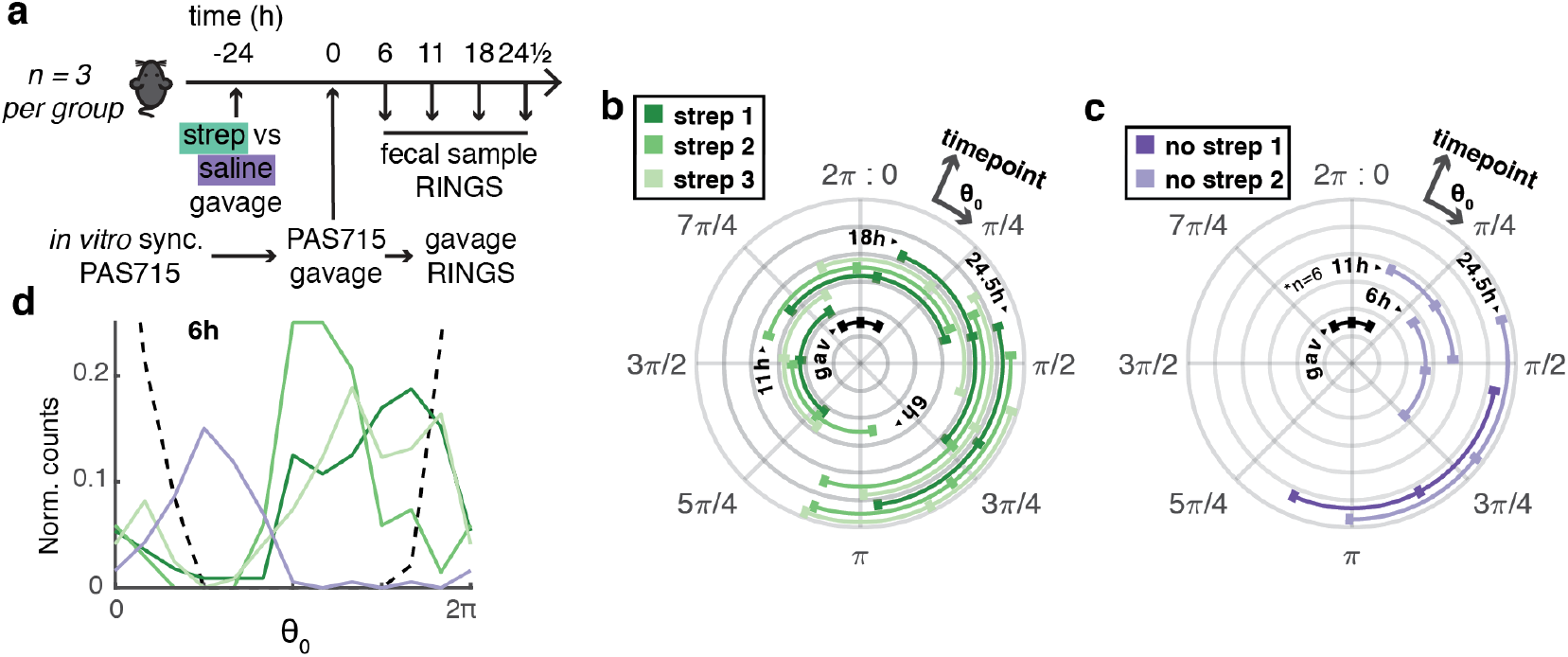
RINGS analysis measures growth variation in the mouse gut. **a.** IPTG *in vitro* synchronized PAS715 bacteria were delivered to mice (n=3 per group) that were provided saline ± 5mg streptomycin by oral gavage 24-hours prior. Samples of the gavaged bacteria and fecal samples collected in the subsequent ~24-hours were grown on selective bacterial plates and RINGS analysis was undertaken to measure repressilator phase progression during transit of the mouse gut. **b-c.** Repressilator phase progression throughout the experiment in **b.** streptomycin treated and **c.** untreated mice. Graphs show circular mean ± angular deviation for colonies from each mouse. Growth corresponds to a clockwise phase shift and each ring corresponds to a discrete timepoint taken. Bacteria were successfully obtained from only 2/3 untreated mice and at low numbers at some timepoints (indicted with *). **d.** Histogram comparison of bacterial phase distribution of gavage (black dotted) with streptomycin treated (green), and untreated (purple), samples from feces collected 6-hours after gavage. Data represent normalized counts from π/6-width bins of phase.

To further test the utility of the repressilator within different bacterial species, IPTG synchronized *E. coli* PAS715 and *S.* Typhimurium PAS716 bacteria were delivered by oral gavage to mice that had been treated 24 hours earlier with streptomycin (5 mg/mouse by oral gavage) (Fig 4a). RINGS analysis was then performed on fecal samples (Fig 4b-e). The growth of each species was quantified using the previously calculated repressilator 2.0 period (Fig 2d-e), with both undergoing similar average growth during the experiment (PAS715: 13.3 ± 0.9 and PAS716: 14.3 ± 0.7 gen at 20.5h post gavage) (Fig 4f). RINGS-estimated growth was most rapid immediately following gavage (Fig 4f), a finding that was further confirmed by Peak-to-Trough Ratio (PTR), which estimates the average instantaneous growth rate of a population from metagenomic sequencing data (6h: 1.72 ± 0.08, n=4; >16h: 1.44 ± 0.08, n = 9; Table 2)^18^. These dynamics are consistent with a model by which rapid replication can occur within the niche made available by streptomycin treatment until resource limitation and recovery of the normal microbiota restricts growth.

**Figure 4:**
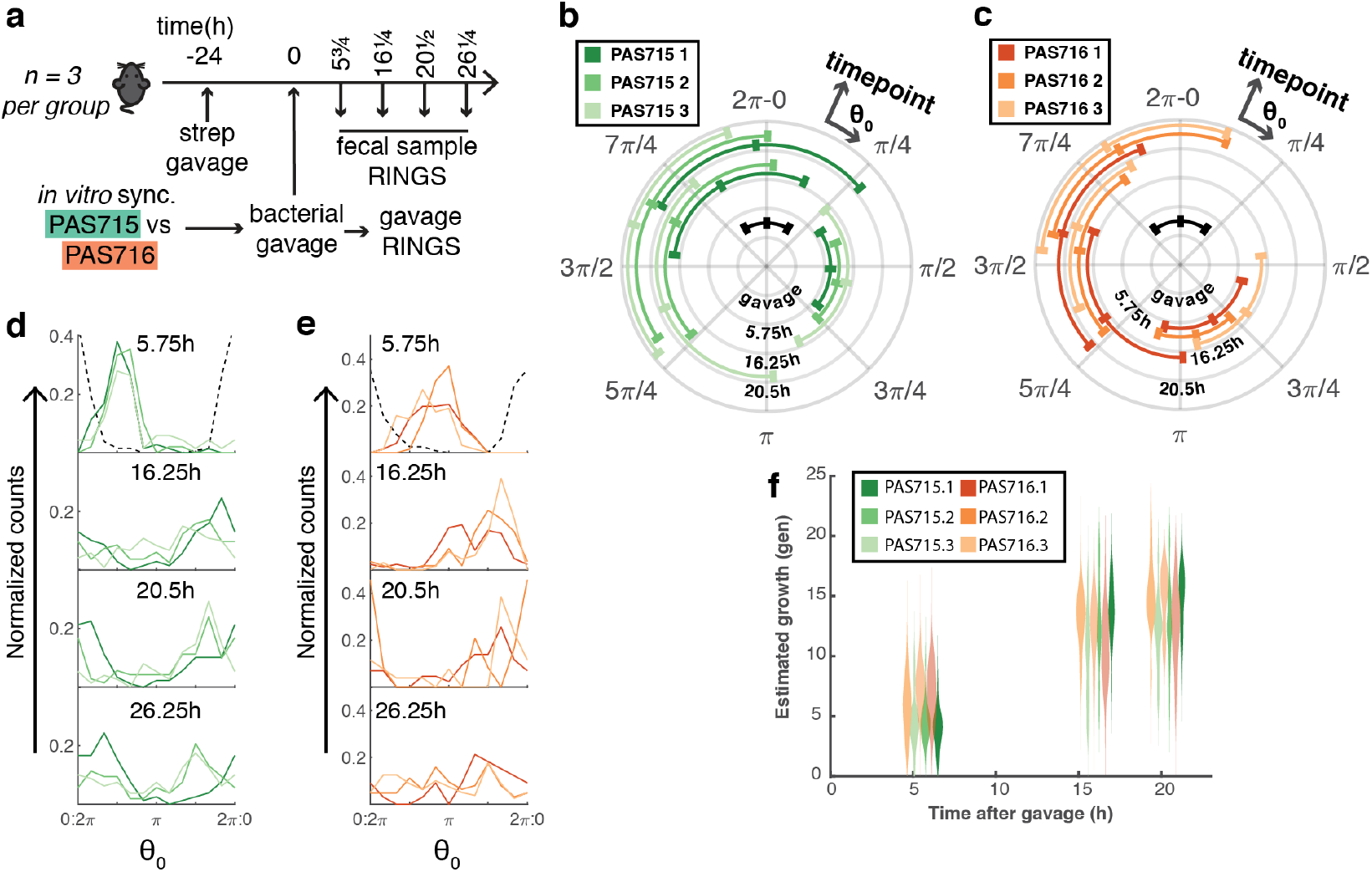
Repressilator 2.0 demonstrates similar behavior between strains. **a.** IPTG *in vitro* synchronized PAS715 *(E. coli*) or PAS716 *(S.* Typhimurium) bacteria were delivered to mice (n=3 per group) that were provided 5mg streptomycin by oral gavage 24-hours prior. Samples of the gavaged bacteria and fecal samples collected in the subsequent ~24-hours were grown on selective bacterial plates and RINGS analysis was undertaken to measure repressilator phase progression during growth within the mouse gut. **b.** PAS715 (green) and **c.** PAS716 (orange) bacterial repressilator phase progression throughout the experiment. Graphs show circular mean ± angular deviation for colonies from each mouse. Growth corresponds to a clockwise phase shift and each ring corresponds to a discrete timepoint taken, **d-e.** Histogram comparison of bacterial phase distributions for **d.** PAS715 and **e.** PAS716 populations from feces collected at each timepoint. 5.75h timepoints are compared to phase distribution of the gavage sample (black dotted line). Data represent normalized counts from π/6-width bins of phase, **f.** Repressilator estimated growth of PAS715 (green) and PAS716 (orange) over time. Graph shows probability density of growth for populations retrieved from individual mice.

**Table 2:** Peak-to-Trough ratio values as calculated from analysis of metagenomic samples sequenced by Illumina sequencing. Higher values correspond to higher estimated growth rates.

| **Bacterial strain** | **mouse** | **Peak-to-Trough Ratio** |  |  |  |
| --- | --- | --- | --- | --- | --- |
|  |  | **Pre-admin** | **5.75h** | **16.25h** | **20.5h** |
| PAS715 | 1 | <5% map | 1.65 | 1.43 | 1.46 |
|  | 3 | <5% map | 1.75 | 1.40 | 1.52 |
| PAS716 | 1 | <5% map | N/A | 1.55 | N/A |
|  | 2 | <5% map | 1.83 | 1.43 | N/A |
|  | 3 | <5% map | 1.66 | 1.36 | N/A |

### The repressilator 2.0 can sense perturbations in the gut over long time periods

To test the ability to control the repressilator 2.0 circuit and re-gain synchronicity of the bacterial populations growing within the mouse gut, mice carrying asynchronous PAS715 bacteria were provided IPTG (10mM) or aTc (0.1 mg/mL) in drinking water overnight for ~17 hours (Fig 5a). RINGS analysis of bacteria plated from fecal samples in pre- and post-synchronized mice clearly demonstrated the ability to synchronize the repressilator within the mouse gut (Fig 5b). To assess the longevity of IPTG’s synchronizing activity in the gut, RINGS analysis was performed on PAS715 fecal samples taken 5, 10 and 28 hours after IPTG overnight provision and removal (Supplementary Fig 4a), with phase only progressing in 28-hour post-synchronization samples (Supplementary Fig 4).

**Figure 5:**
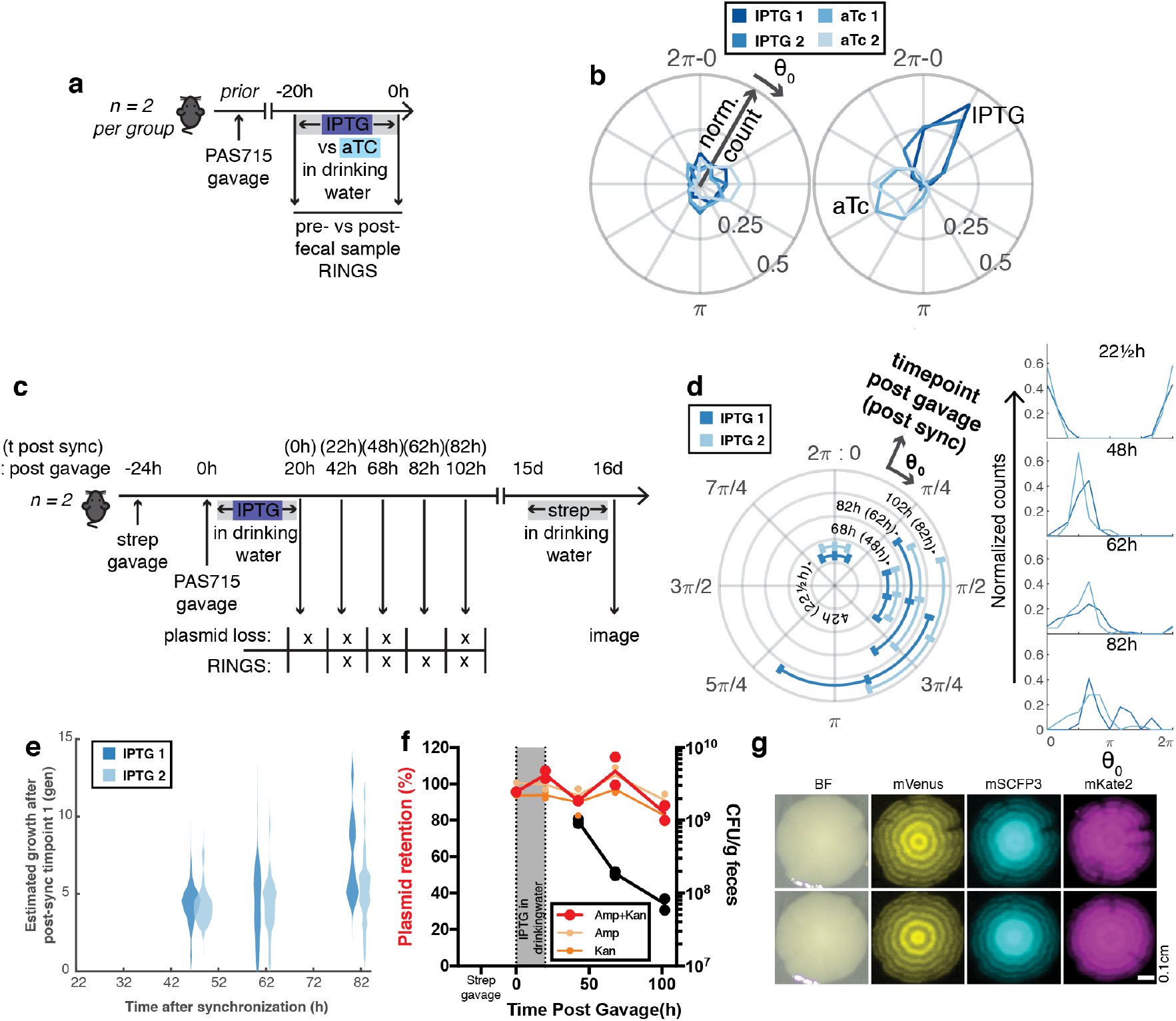
The repressilator 2.0 is stable over time and can be controlled within the mouse gut. **a.** Mice (n= 2 per group) carrying asynchronous PAS715 were provided IPTG (10mM) or aTc (O.1mg/mL) with 5% sucrose in drinking water for ~17h overnight with fecal samples taken before and after provision of the inducer. **b.** RINGS analysis demonstrated the progression from asynchronous (left) to synchronous (right), with IPTG (darker blue) and aTc (lighter blue) synchronizing cultures to different phases of the repressilator. **c.** Repressilator *in situ* synchronization allows RINGS analysis later in colonization. Mice (n=2) were provided asynchronous PAS715, followed by overnight ~20h provision of IPTG (10mM) in 5% sucrose drinking water. Fecal samples were then collected ~ every 24-hours and plated on selective plates to assay for repressilator and sponge plasmid retention, or RINGS analysis to follow repressilator phase. After 15 days, provision of streptomycin (0.5g/L) in 5% sucrose drinking water overnight selected remaining PAS715 bacteria to allow isolation of colonies that had been present in the gut over at 16-day period. **f**. RINGS analysis of phase progression 42-102 hours after gavage. Circular graph (left) shows circular mean ± angular deviation for colonies from each mouse. Growth corresponds to a clockwise phase shift and each ring corresponds to a discrete timepoint taken. Linear histograms (right) compare bacterial phase (Θ_0_) distributions throughout the experiment. Data represent normalized counts from π/6-width bins of phase. **g**. RINGS-estimated growth following IPTG removal from the drinking water. Graph shows probability density function of the population retrieved from individual mice. **h**. Plasmid retention (colored samples, left axis) and bacterial abundance (black samples, right axis) data from plating on differentially selective plates. Graph shows individual values and mean line.

The repressilator is stable and remains functional over long periods in the mouse gut. We delivered unsynchronized PAS715 bacteria to mice (n=2) that had previously been treated with streptomycin (Fig 5c). Mice were immediately provided IPTG for ~20 hours within their drinking water to synchronize the repressilator *in situ*, after which they were returned to normal drinking water. Following a 24-hour period to allow repressilator progression to re-commence, RINGS analysis was performed on fecal samples plated daily to estimate bacterial growth rate. The repressilator 2.0 remained relatively synchronous within the gut without extensive progression through the oscillator’s period for at least 80 hours following removal of IPTG from the drinking water (Fig 5d), which corresponded to 5-10 generations over a 60-hour period (Fig 5e).

We demonstrated the stability of the repressilator circuitry by plating of fecal samples on different combinations of selective medium in order to identify plasmid loss events within the bacteria (Fig 5c). Growth on streptomycin (all PAS715) was compared to streptomycin + carbenicillin (retention of repressilator plasmid), streptomycin + kanamycin (retention of sponge plasmid), and streptomycin + carbenicillin + kanamycin (retention of both plasmids). Plasmid retention of both the repressilator and sponge plasmids remained high throughout the experiment (80-88% retention at 102h post gavage), indicating that the repressilator circuit does not provide sufficient burden to unduly affect bacterial growth in the gut (Fig 5f). Furthermore, 15 days post gavage, mice were provided a fresh dose of streptomycin (0.5 g/L in drinking water overnight) in order to allow re-expansion of PAS715 bacteria remaining within the gut. We were able to isolate functional PAS175 colonies 16 days after first entering the gut (Fig 5g), demonstrating the potential for this synthetic circuit to maintain oscillatory gene expression over considerable periods of time in a competitive environment.

### The repressilator measures disease perturbations within the gut

We used the RINGS method to examine the effects of inflammation on PAS715 colonization using the Dextran Sulfate Sodium (DSS) inflammation model^19^. Inflammation, including DSS treatment, is known to promote colonization and increase growth of *Enterobacteriaceae* within the gut^20^. Inflammation group mice (n=3) were given DSS (3.5% in drinking water) for 3 days, before IPTG synchronized PAS715 were provided by oral gavage (Fig 6a). Repressilator 2.0 period length was unperturbed during mother machine growth on cecum fluid extracted from DSS treated mice (Table 1). Bacterial growth was therefore measured by RINGS analysis on fecal samples over the following 24 hours. Bacteria in DSS treated and untreated (no strep) controls showed similar RINGS phase after 5 hours (Fig 6b), however, the populations retrieved from DSS treated mice >19 hours after gavage all showed strong indications of subpopulations growing at different rates (Fig 6b). Analysis of the angular deviation at each timepoint showed that bacterial populations in DSS treated mice had the most extreme loss of synchrony observed across all experiments (Fig 6c).

**Figure 6:**
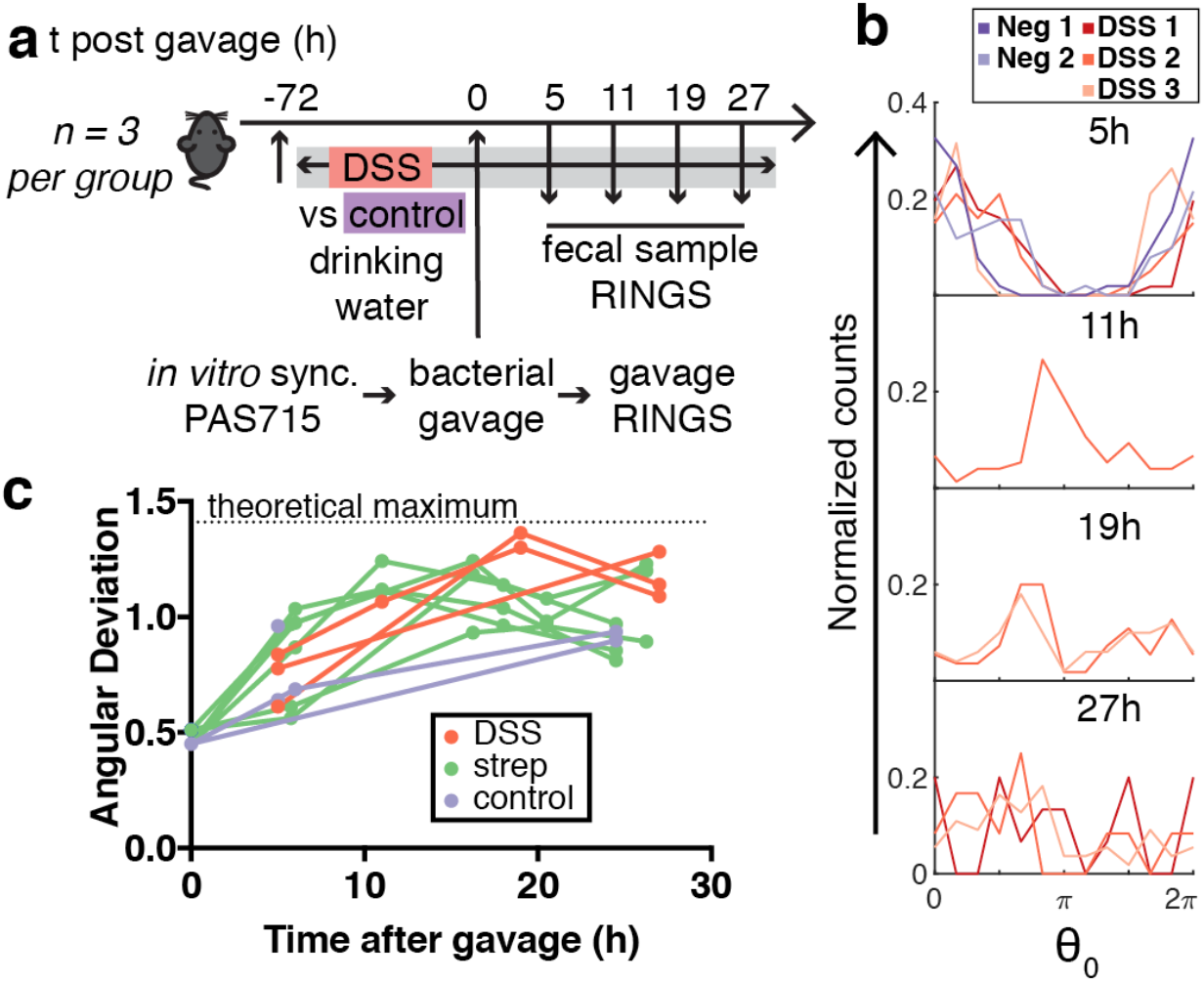
Bacterial growth in the inflamed gut is variable. **a.** IPTG *in vitro* synchronized PAS715 were delivered to mice (n=3 per group), that had previously been fed dextran sulfate sodium (DSS) or normal drinking water for 3 days prior. Samples of the gavaged bacteria and fecal samples collected in the subsequent ~24-hours were grown on selective bacterial plates and RINGS analysis was undertaken to measure repressilator phase progression during growth within the inflamed mouse gut. **b.** Histogram comparison of bacterial phase distribution of PAS715 in DSS treated (red) or and untreated (purple) samples from feces collected each timepoint. Data represent normalized counts from π/6-width bins of phase (θ_0_) **c.** Plots of angular deviation over time for PAS715 mouse experiments (Figs 3–5) in DSS (red), streptomycin (green) or untreated (purple) mice.

## Discussion

Here we describe the activity of a synthetic gene oscillator, the repressilator 2.0, in the complex environs of the mammalian gut. We develop an image processing and analysis pipeline, RINGS, to follow repressilator activity on a single-cell level and thereby also understand growth dynamics at various stages of the colonization process. We demonstrate robust repressilator 2.0 functionality, control and circuit stability within the mouse gut for several weeks. Our results show that single-cell measurements are particularly important in disease conditions, such as inflammation, when variability of growth between niches is accentuated and thus methods that can only measure a population-wide average do not adequately represent the population’s diversity.

RINGS analysis estimated generation times of *E. coli* MG1655 progressively increasing from ~35 minutes to ~10h as the native microbiota recovered from a single dose of streptomycin, which can be used to create an initially favorable growth niche for *E. coli* within the gut. These numbers are comparable to those calculated using previous methods for growth measurement, recent examples of which include PTR for instantaneous rate measurements from metagenomic sequencing data^18^, a synthetic particle which allows mark-and-recapture analysis over ~14 generations^21^, and fluorescent protein dilution which is capable of estimating up to ~10 divisions^22,23^.

Previous studies provide a model, the ‘restaurant hypothesis’, for *E. coli* colonization of the mouse gut, whereby strains colonize or not depending on the availability of a unique and compatible ‘nutrient-niche’, with competition occurring as planktonic cells in the gut lumen^24^, but replication preferentially occurring in multispecies biofilms within the mucus layer^25^. The spatial segregation of independent biofilms may allow competing strains to co-exist with reduced competition if the initial colonization resistance is overcome. Our finding that similar growth variability across the recovered population occurred in mice with normal (untreated) and reduced (streptomycin treated) colonization resistance, despite differences in overall bacterial load of 4-8 orders of magnitude, supports this model, suggesting that in rare cases where bacteria overcame active colonization resistance they grew in a uniform manner. Our findings of a more rapid repressilator desynchronization under inflammatory conditions can also be interpreted to suggest that spatial variation in inflammation leads to increased variability of growth environments that are not uniformly experienced by the population, nor overcome by mixing between niches.

Of particular note, if we look at the fluorescent reporter proteins as placeholders for any given transcriptionally controlled protein or mRNA, the repressilator 2.0 is a prime candidate for periodic expression control for use in complex settings. This opens the door for translational advances such as periodic therapeutic delivery^26^ or growth-linked control of higher-order synthetic circuits such as counters^27^, or memory circuits^28–30^. A recent demonstration of pulsatile lysis and toxin delivery by engineered bacteria within tumors, achieved through quorum sensing at high bacterial concentration, improved survival in a colon metastasis model^15^. The repressilator’s independence from external signals would allow for circuit function in a variety of environments, which is of particular utility in the gut where relative abundance of strains can vary widely on spatial and temporal scales, and between individuals. To this end, the ability to re-synchronize the repressilator 2.0 *in situ* within the gut, and its stability over extended periods of growth without antibiotic selection are particularly exciting.

In sum, analysis of the repressilator at single cell level can report on complex bacterial growth behaviors over time, and in particular, during disease conditions within the mammalian gut. In doing so, this work demonstrates the potential for one of the circuits that first stimulated the field of synthetic biology to revolutionize how we control gene expression within the gut, both for research and for therapeutic purposes.

## Materials and Methods

### Strains, plasmids and bacterial culturing

Details of plasmids (Supplementary Table 1) and strains (Supplementary Table 2) used in this study are provided.

The repressilator 2.0 plasmid variant with three fluorescent reporters (pLPT234) was constructed by isothermal assembly^31^ by combining PCR products from previously published degradation-tag free repressor genes – Tn10 transposon derived *tetR*, bacteriophage λ derived *cI* and *E. coli* lactose operon derived *lacI* (pLPT119) – and triple fluorescent reporters – PR-mKate2 (including the first 11 amino acids of mCherry for improved translation efficiency as previously published^8^), PLtet01–mVenus, PLlac01– mSCFP3(pLPT107)^8^. Sponge plasmid variants were constructed as reported previously^8^. Plasmids were isolated by miniprep (Qiagen) and were routinely transferred to new strains as a mixture by electroporation.

To avoid interruption of the repressilator and ensure clear ring development within colonies, *lacI* and *motA* genes were knocked out of *E. coli* strains used before repressilator plasmids were transferred. For both genes, FRT-flanked gene disruption constructs were transferred by P1 *vir* transduction^32^ from the relevant Keio collection strains^33^. Kanamycin resistance genes were then removed by electroporation and subsequent curing at 43°C of pCP20^34^ Similarly, streptomycin resistance based on a *rpsL* lys42arg mutation was transferred by P1 *vir* from a previously generated *E. coli* MG1655 strain^29^. Resistance to streptomycin was evolved in *S.* Typhimurium LT2 by serial passage in liquid culture with increasing concentrations of streptomycin sulfate (Sigma) (50,100, 200, 300 μg/mL). For PAS718, mKate2 (including the first 11 amino acids of mCherry for improved translation efficiency as previously published^8^) driven under the Prna1 constitutive promoter was inserted into the genome using a Tn7 transposon^35^, and acted as a constitutive fluorescent marker for cell segmentation in mother machine experiments.

Bacteria were routinely cultured in Luria broth (LB) supplemented with 300μg/mL streptomycin (Sigma), 100μg/mL carbenicillin or ampicillin (Sigma) and 50μg/mL kanamycin (Sigma). For plating, bacteria were grown on selective LB agar plates supplemented with 100μg/mL carbenicillin (Sigma) and 50μg/mL kanamycin (Sigma).

To synchronize the phase of the repressilator across the population, bacteria were grown overnight in the presence of 1mM Isopropyl β-D-1-thiogalactopyranoside (IPTG) (Sigma) or 100nm anhydrotetracycline (aTc) (Sigma). Bacteria were backdiluted in fresh inducer-supplemented media by at least 1:20 to allow resumption of active growth in the presence of inducer, before being washed and utilized for downstream experiments.

### Colony imaging

Fluorescent and white light images of colonies were imaged using a custom-software controlled Canon T3i digital single lens reflex (DSLR) camera with a Canon EF-S 60 mm USM lens, combined with LEDs and filters for excitation and a Starlight express emission filter wheel (CFP: 440-460nm LED with 436/20 EX and 480/40 EM filters; YFP: 490-515nm LED with 500/20 EX and 530/20 EM filter; RFP: 588-592 LED with 572/35 EX and 645/75 EM filter; white: 3500-4500K LED)^36^. Images were taken at an aperture of f/2.8 and IS0200. Exposure times were typically between 0.05 and 2s as experimentally determined to maximize dynamic range.

### RINGS method for determining phase offset in repressilator colonies

We extract the relative phase offset of each colony by fitting a generative model to the pattern of oscillations. This approach has the benefit of making use of all the information captured in the multi-channel images. We explored a family of generative models, based on the simple form:

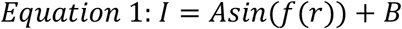

where A and B are the amplitude and offset of a sinusoidal oscillator, and the function, f (r), represents the radial phase profile of the colony. The form of f (r) is derived in *Growth Model*, using a simple model for the growth of the colonies.

### Growth Model

The phase profile, *f(r)*, is related to the profile of generation number, *g(r)*, as follows:

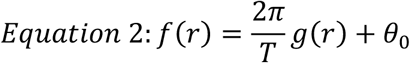

where T is the period of the oscillator in generations (≈ 15.5) and Θ_0_ is the instantaneous phase of the oscillator at the time of plating. Thus, we derive the profile of generation number with respect to radius. We assume that growth is initially exponential, and then becomes restricted to an annulus with thickness D (≈ 30μm at the edge of the colony^37^).

#### Phase I: Exponential growth

Assuming uniform packing of bacteria in the colony (for both phases of growth), the generation number profile during exponential growth can be derived simply:

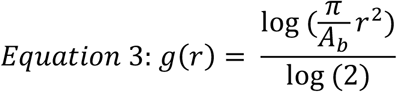

where A_b_ is the area of a single bacterium

#### Phase II: Annular Growth

Here we model the growth of the colony with each successive generation. With each generation the radius of the colony increases as follows:

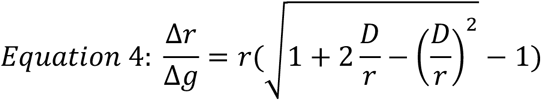

Integrating this over the colony, we obtain the following expression for the generation number profile:

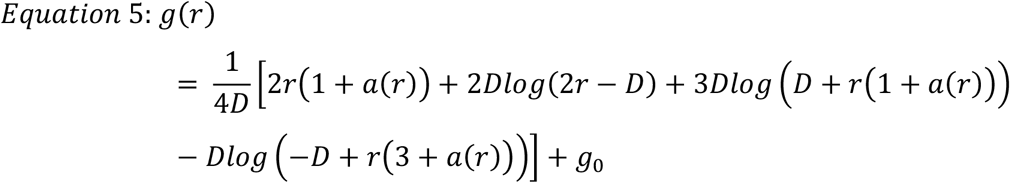

g_o_ is the generation number at the transition between the exponential and annular growth.

Combining the two growth models yields a continuous growth curve (Supplementary Fig 5). We set the transition between the exponential and annular growth at r = D, with D= 30μm. Notice that beyond r ~ 100μm the phase profile is approximately linear, and can be described by:

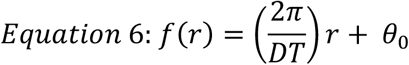

We checked this result by measuring the slope for the phase profile directly from images of colonies. The distance between two peaks in any given colony is approximately 490μm, yielding a phase profile with slope 0.0128 rad/μm.

Using an approximate value of 15.5 generations per cycle, equation 7 predicts an annulus size, D ≈ 32μm, consistent with the ≈30μm growth annulus reported in previous studies^37^.

Thus, we model the phase profile of a bacterial colony using Equation 1 with a linear phase profile, described by Equation 6.

### Generative Model Fitting

Images were initially pre-processed as described above, to crop centered colonies from whole-plate images, and normalized for global changes in intensity, before fitting the model described above.

### Preprocessing

Centered colonies were cropped from whole-plate images using FIJI^38^. Color images from single fluorescent exposures in CFP and YFP channels were imported, and the relevant spectral component image utilized for downstream processing (CFP – blue channel, YFP – Green channel). Colonies were identified by autothresholding (Yen method) of binarized CFP images. Separate YFP and CFP images were cropped and exported, centered on the center of mass of each identified colony. Prior to final analyses, colony images were visually inspected and any severely malformed or doublet colonies were removed from the datasets to avoid spurious fitting. Colony removal was performed blinded to Θ_0_ fit values. Average projections were taken through all colonies identified in a given population.

### Model fitting

We observed that in each colony, the expression of YFP/CFP decayed dramatically with increasing radius. To compensate for this decay, the images were masked at radius of r_max_ =1.3mm (r_max_ =1.1mm for *S*. Typhimurium colonies which were commonly smaller) and the radial intensity profile was fit to a second order polynomial. This smooth polynomial decay, chosen as an even order with negative coefficient to ensure this process didn’t removed the oscillatory patter, was then used to normalize the raw image in a pixel-wise fashion (Supplementary Fig 6). We also explored normalization with a fourth order polynomial, but this was deemed less appropriate. For multi-channel images, the polynomial functions shared parameters for the center of the colony, but each channel was fit to its own decay profile.

To fit the oscillatory pattern of a single colony, we use Equation 1 with a linear phase profile. However, we allow for both the slope and offset of the phase profile to be fit (Equation 6), as well as the amplitude and offset of the intensity (Equation 1). The slope for the phase profile is initialized to an expected value that varies based on strain, and the phase offset is initialized to zero. We tested optionally masking the central region of the colony, to account for the early exponential growth phase, which we don’t attempt to model, however found no benefit to fit based on this. Fitting is done in Matlab using the Levenberg-Marquardt (LM) algorithm (Supplementary Fig 6).

The fitting routine returns the center position of the colony, the amplitude and offset of the oscillations, as well as the slope and offset of the phase profile. The phase offset is the property of interest. However, fitting the radial expression profile is challenging due to numerous local minima in the error surface. Thus, we filter out colonies for which the slope of the phase profile, or the inferred center position of the colony is outside of an accepted range. We also explored the use of robust error functions, to minimize the effect of asymmetric bright patches of expression in the colony, as well as weighted nonlinear regression to compensate for the increasing number of pixels at increasing radius. However, neither of these approaches greatly improved upon the LM algorithm, which we used by default. We also explored fitting phase profiles with higher order polynomials; however, we found this to be very unstable.

Fitting was improved by simultaneous regression in multi-channel images. In this case, the phase offset of the two channels was fixed to a constant value. We found that YFP oscillations lead CFP by a strain-specific value, which can be visualized by independently fitting YFP and CFP images from the same population (Supplementary Fig 7). Thus, we regressed the slope and offset of the phase profile using both channels, with a fixed phase difference between the two channels, as calculated for each strain. See Equation 7.

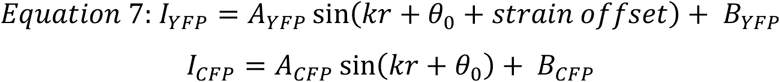

### Testing the method

We tested the performance of this method by applying it *in vitro* timecourse data where colony counts could be used to provide a known generation shift between each timepoint. Parameter sweeps on key parameters expected to vary between strains, particularly the size of colony mask (‘rMax’), the expected slope of the phase profile (‘expectedSlope’), the variation in slope between colonies (‘slopeTol’), the distance between each ring, and the offset between CFP and YFP rings (‘colorPhaseShift’). An example of this process is provided in Supplementary Fig 8, which demonstrated that the method was largely insensitive to specific parameter values. Optimal parameters for each strain used in the study are as follows: LPT320: ‘rMax’:1.3, ‘expectedSlope’: 0.39, ‘slopeTol’: 0.3, ‘colorPhaseShift’: 1.5. PAS715: ‘rMax’:1.3, ‘expectedSlope’: 0.34, ‘slopeTol’: 0.4, ‘colorPhaseShift’: 0.9. PAS716: ‘rMax’: 1.1, ‘expectedSlope’: 0.43, ‘slopeTol’: 0.4, ‘colorPhaseShift’: 1.0.

### Modulation for elapsed phase measurements

Modulation of phase measurements was predominantly undertaken using custom scripts in Matlab versions 2016a-2018a. In order to estimate total phase elapsed during experiments in which the repressilator progressed more than a single period, we made the assumption that repressilator phases across the population fell within ± π of the circular mean (i.e. we assumed that all colonies were within 15 generations elapsed of each other). We deemed this appropriate where the population remained relatively coherent around the mean but refrained from this form of analysis where signiifcant portions of the population could by mis-attributed as faster/slower growing or where no clear population mean was apparent.

For calculations, all datapoints were first normalized by modulo 2π of the circular mean at timepoint zero. Θ_0_ datapoints were then adjusted by multiples of 2π based on their modulo 2π value, according to the scenarios depicted in Supplementary Fig 9. Subsequent calculations treated the resultant values as linear datasets.

To calculate probability density functions (PDFs), initial synchronized repressilator 2.0 populations (t_sync_) were assumed to be normally distributed, which was appropriate based on visual inspection of Q-Q plots. Accordingly, individual datapoints were calculated to have a PDF for phase elapsed from t_sync_ as a normal distribution centered on Θ_0_ (since μ_θ_(t_sync_) = 0) with standard deviation σ_tsync_. Subsequent datasets were not assumed to follow a normal distribution, but instead PDFs were calculated as evenly weighted sums of all PDFs of individual datapoints. Conversion from phase to generations was calculated based on values determined using the slope of linear regressions calculated between RINGS analysis and CFU counts (Fig 2d-e).

## Growth in the mother machine

We performed time-lapse single-cell measurements to monitor and compare the dynamics of repressilator circuits in rich defined medium (EZ Rich Defined Medium; Teknova) and in cecum-contents. Cecum contents were derived from fresh or frozen mouse cecum, diluted in PBS. 2-4 FVB mice (Charles River) of varied age were sacrificed, their cecum dissected, and the cecum contents were extracted by scraping and washing off the tissue with PBS. When frozen, cecum contents were placed at −80°C. The contents were then thawed, if necessary, and diluted in PBS, spun briefly in a benchtop centrifuge (13,000g) to remove large particulate matter and the supernatant removed and saved. Cecum matter was then washed a second time, centrifuged, and the supernatant pooled with the first wash. Pooled supernatant was centrifuged a second time, then passed sequentially through a 20 μm syringe filter (Millex-AP glass fiber 25mm syringe filter) and then a 5 μm syringe filter (Pall Acrodisc 32mm syringe filter with 5μm Supor^®^ membrane). The filtered fluid was then used as an input for the mother machine.

The filtered fluid from the cecum content was loaded in 10 mL syringes and pumped through mother-machine devices^17^ using a Higher Pressure Programmable Syringe Pump (NE1000). To flow the cecum through the device, we used a mother-machine design where the dimensions of the flow channel has been optimized for flowing dense cultures and maintaining low pressure to allow low fluid flow-rates (Bakshi, S., Leoncini, E., Baker, C., Canas-Duarte, S., Okumus, B. and Paulsson, J., unpublished data). This particular design consists of flow channels with 150 μm width and 45 μm height. Cells are loaded in narrow trenches (25 μm long, 1.5 μm wide and 1.3 μm height) that are placed in orthogonal direction to the flow channels. The details of the device preparation from the wafer and loading cells is described previously^8^.

Prior to flowing the content of the cecum, we loaded the trenches with E. coli containing the repressilator circuit. In some experiments *E. coli* PAS718, constitutively expressing mKate2, was used, with the mKate2 acting as a marker to segment single cells and to extract intensities and cell-growth estimates more effectively. We used a slow flow-settings (5 μL/min) to ensure we can observe the dynamics of the cells for prolonged periods (>10 h) with the small volume of cecum content (~ 3-5 mL). This ensured that we observed at least two peaks for repressilator signals in individual channels (YFP or CFP) for a majority of the cells, which is necessary to calculate the period of oscillation. To minimize phototoxicity the frequency of imaging was kept at 6 min/frame. This gives about 4-5 snapshots per generation time, which is enough to get a good estimate of growth-rate and also a smooth intensity time-series. For *E. coli* PAS715 experiments imaging occurred at 10min/frame.

Images were acquired using a Nikon Ti inverted microscope equipped with a temperature-controlled incubator (OKO lab), a sCMOS camera (ANDOR), a 40X Plan Apo air objective (NA 0.95, Nikon), an automated xy-stage (Nikon) and light engine LED excitation source (Lumencor SpectraX). All experiments were performed at 37°C. Microscope control was done with Nikon Elements software. We acquired time-lapse data from 40 fields of view, which allows us to track approximately 2000 cells simultaneously. The acquired data was analyzed using a hybrid analysis platform written in the Paulsson lab. In brief, images were segmented using a custom-designed FIJI plugin and then the extracted data was further processed to track cells in a custom-designed MATLAB script. Time-series of cell-size data calculated from the segmented images were used to compute generation times. The time-series of YFP channel intensity was used to calculate the period of repressilator in the cecum content.

## *In vivo* bacterial growth testing

The Harvard Medical School Animal Care and Use Committee approved all animal study protocols.

Female C57Bl/6 (Jackson Laboratory) mice of 8-14 weeks, (including >2 weeks acclimatization to the HMS mouse facility), were used. Mice were routinely randomized between treatment groups in advance of experiments, where relevant. Mice were fed on a lactose-free chow (Envigo Teklad Global 18% Protein Rodent Diet) for at least 1 week prior to provision of repressilator bacteria to avoid interruption of the repressilator by lactose, however, subsequent *in vitro* experiments suggested this was not a concern (Supplementary Fig 10). Streptomycin treated mice were administered 5mg USP-grade streptomycin sulfate (Gold Biotechnology) in 100μL sterile PBS by oral gavage, or where specifically stated 0.5g/L in drinking water supplemented with 5% sucrose overnight. PAS715 *E. coli* MG1655 bacteria or PAS716 *S.* Typhimurium LT2 bacteria were prepared for administration by pelleting from culture, washing and dilution in sterile PBS before provision to mice as a 100μL oral gavage.

*In situ* synchronization was achieved by provision of USP-grade 10mM IPTG (Sigma) or 0.1mg/mL aTc (Sigma) in drinking water supplemented with 5% sucrose overnight.

Fresh fecal samples were collected by temporarily removing mice to small containers until at least 2-3 fecal pellets were produced. Fecal pellets were homogenized in PBS at 50 or 100mg/mL in sterile PBS by vortexing for ~5min in 1.5mL Eppendorf tubes. To remove large debris, homogenized feces was then centrifuged either by briefly pulsing (~1sec) on a benchtop minifuge (where CFU counts were not critical), or by centrifugation at 200 rpm (4g) for 20 min (where CFU counts were critical). The supernatant was then serially diluted and cultured on selective agar plates.

### DSS inflammation model

For inflammation experiments, mice were fed Dextran Sulfate Sodium (Colitis Grade, M.W. = 36,000-50,000, MP Biomedicals, LLC) in drinking water supplemented with 5% sucrose for 3 days prior to bacterial administration. DSS water was exchanged every second day, and mice remained exposed throughout the course of the experiment.

## Peak-to-Trough ratio analysis

Peak-to-trough ratios were computed from metagenomic sequencing of mouse fecal samples as previously described^18^. Genomic DNA was extracted from flash-frozen fecal samples using a Qiagen DNEasy Blood & Tissue Kit. Genomic DNA from each sample was prepared for sequencing using a Kapa HyperPlus Kit and sequenced using an Illumina NextSeq at the Bauer Core Facility at Harvard University. Sequencing reads were trimmed for quality using Trimmomatic 0.3 6^39^ and each sample was aligned to the genome of interest using BWA mem 0.7.8^40^. Sequencing coverage was obtained using bedtools 2.27.1^41^. To calculate peak-to-trough ratio, mean coverage was computed for 10 kb bins across the genome and a segmented linear model was fit to the coverage using the R package segmented^42^.

## Statistical analyses

Statistical analyses were performed in MATLAB versions R2017a-2018a (Mathworks) or Prism v6-7 (Graphpad).

Circular statistical tests were performed where relevant, using the CircStat for Matlab toolbox v2012a^43^.

For histograms, Θ_0_ data were separated into π/6-width bins centered on 0 and multiples of π/6. Counts were normalized by the total number of fit colonies in each dataset. Datapoints were plotted at each bin center. For linear histograms, 0 and 2π are plotted as the same value.

Many figures utilize color palettes based on research by Cynthia Brewer^44^

## Acknowledgements

We would like to thank Lorena Lyon for experimental assistance. Next-generation sequencing was undertaken at The Bauer Core Facility at Harvard University. D.T.R. was supported by a Human Frontier Science Program Long-Term Fellowship and an NHMRC/RG Menzies Early Career Fellowship from the Menzies Foundation through the Australian National Health and Medical Research Council. A.A.N. was supported by an NSF Graduate Research Fellowship (DGE1745303). The research was funded by Defense Advanced Research Projects Agency Grants HR0011-15-C-0094 (P.A.S.) and HR0011-16-2-0049 (J.P. and P.A.S.), and the Wyss Institute for Biologically Inspired Engineering.

## Author Contributions

Conceptualization: D.T.R; Methodology: D.T.R, D.L.R., A.D.N., S.B., E.L., J.P., P.A.S.; Software: D.T.R., D.L.R,; Formal Analysis: D.T.R., A.A.V., A.D.N., S.B., E.L.; Investigation: D.T.R., A.A.V., A.D.N., S.B., E.L.; Resources: L.P-T, A.A.V.; Writing – Original Draft: D.T.R., P.A.S.; Writing – Review & editing: all authors; Visualization: D.T.R.; Supervision: D.T.R., J.P., P.A.S.; Project Administration and Funding Acquisition: J.P., P.A.S.

## SUPPLEMENTARY MATERIAL FOR

**Supplementary Figure 1.**
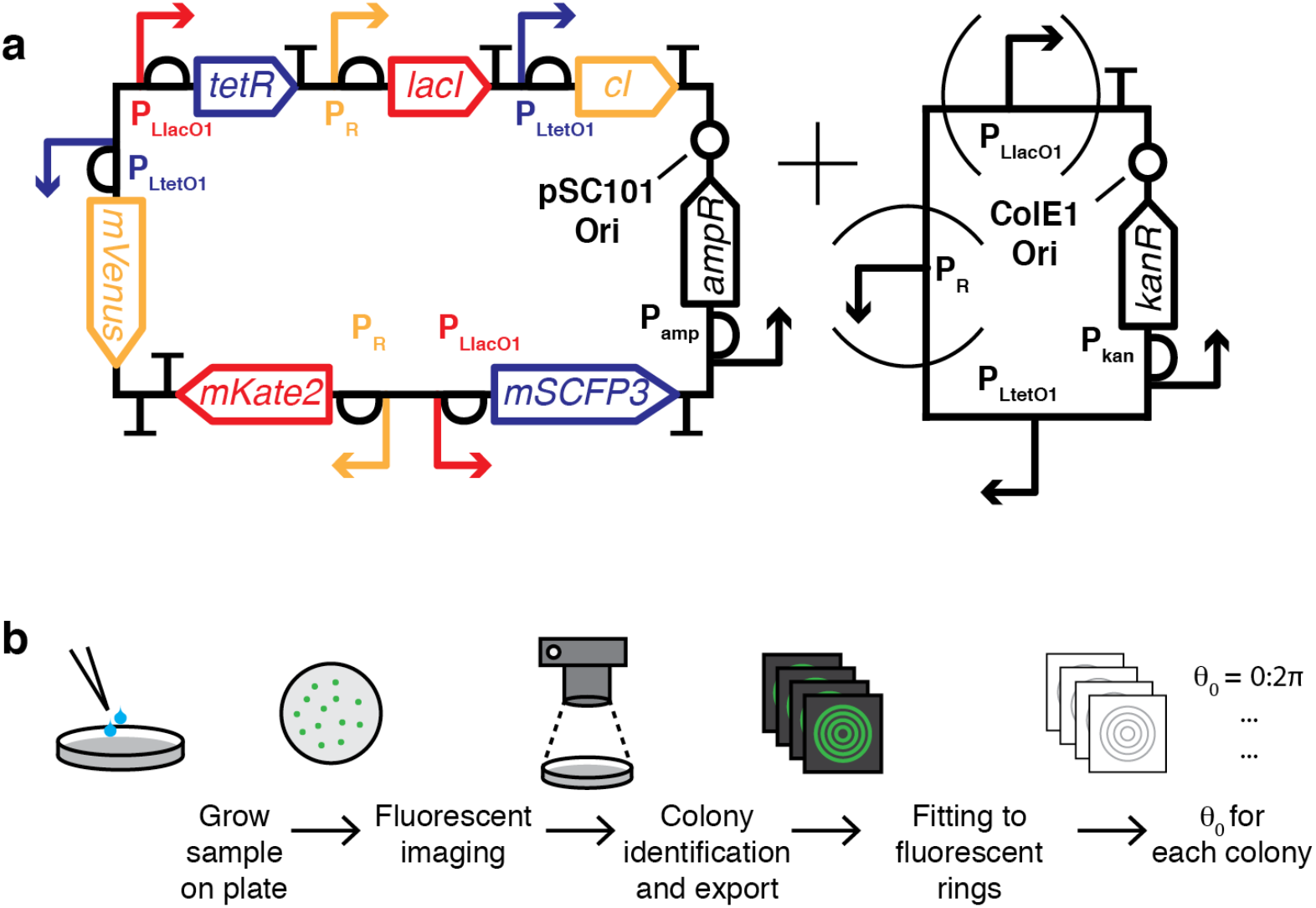
The repressilator 2.0 and RINGS pipeline. **a**. Schematics of the triple-reporter repressilator and sponge plasmids used throughout the study. **b**. The RINGS pipeline consists of plating of a bacterial population on agar plates, followed by imaging, computational identification and export of individual, centered colonies and fitting of a generative model. The fit of ring position is then used to estimate θ_0_ at the time of plating.

**Supplementary Figure 2.**
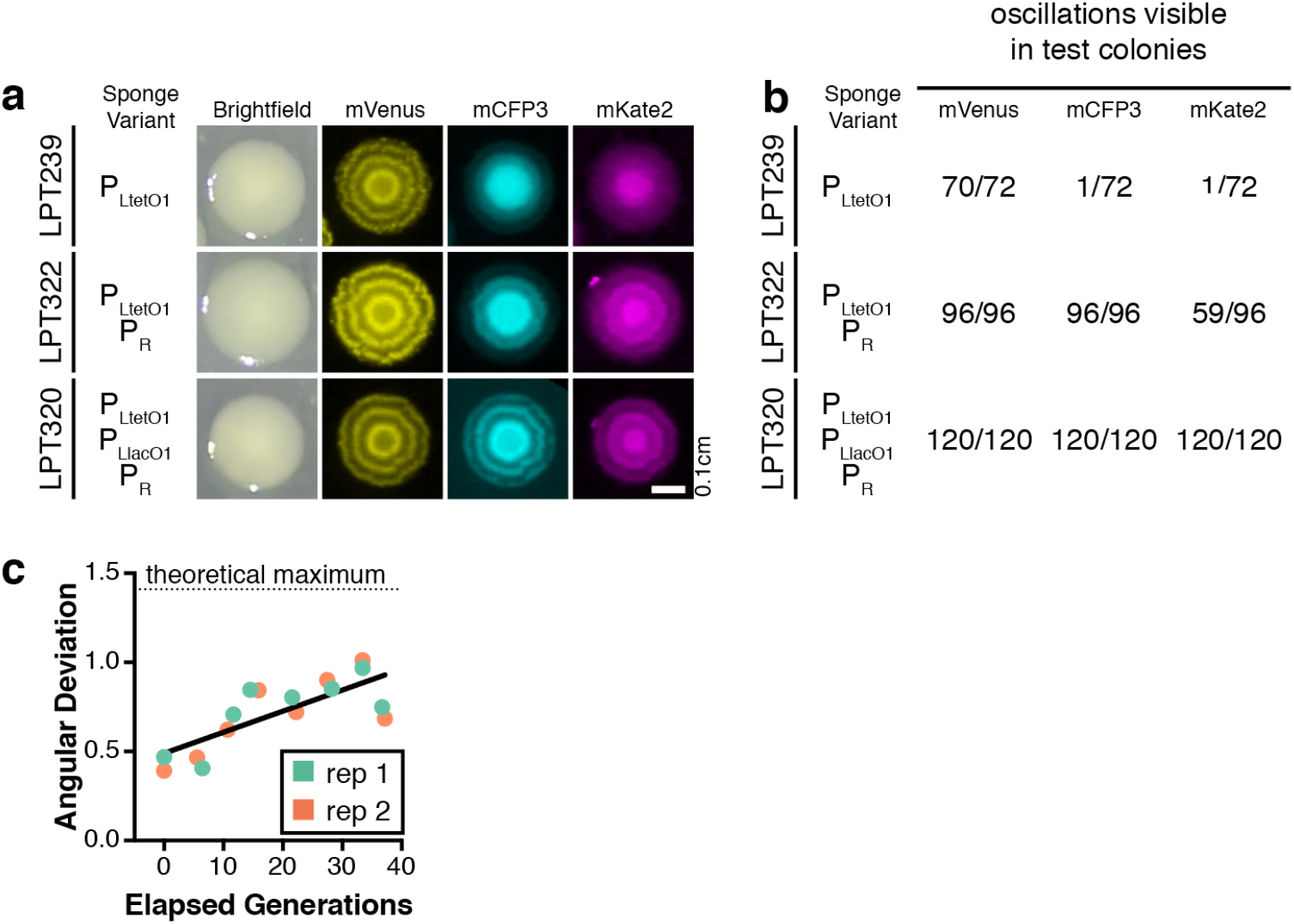
multicolor RINGS analysis and testing. **a.** Fluorescent imaging of colonies formed by *E. coli* MC4100 bacterial carrying the repressilator 2.0 along with sponge plasmid variants – P_LtetO1_ only (LPT239), P_LtetO1_ + P_R_ (LPT322) and P_LtetO1_ + P_R_ + PLlacO1 (LPT320) – showed variability in the consistency of fluorescent rings formed. Scale bar = 0.1cm **b.** Visual inspection of colonies from each strain also showed the clear presence of fluorescent rings differed across each population. **c.** Angular deviation of LPT320 during ~14 hours and ~40 generations of log-phase growth.

**Supplementary Figure 3.**
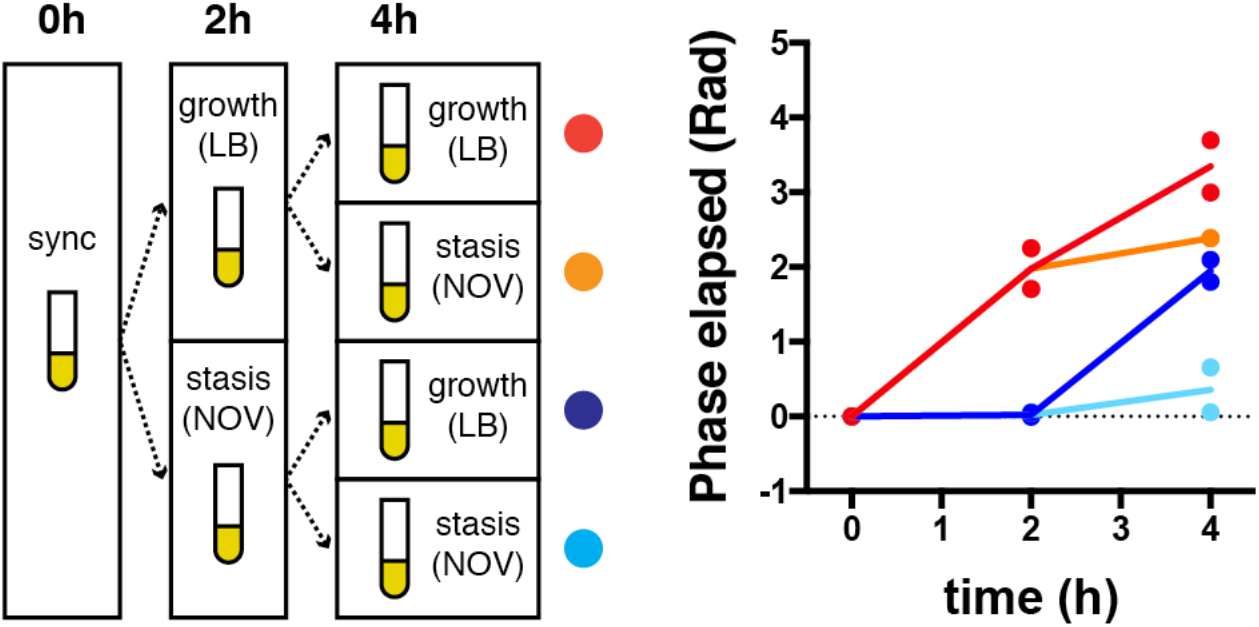
RINGS is unaffected by variable bacterial growth. To test the ability for RINGS analysis to track and integrate growth under variable conditions, IPTG-synchronized PAS715 was back-diluted and grown in liquid culture in the presence or absence of bacteriostatic concentrations of novobiocin. Following two hours of growth, each culture was washed and split again to be grown under the presence or absence of novobiocin, generating four unique combinations of expected growth and non-growth. The phase of populations sampled at 2 and 4 hours were consistent with the expected growth phenotype. In particular, the equivalence of the two populations exposed to one period with and one period without novobiocin, irrespective of order, demonstrates the ability to measure absolute growth during periods of variable growth rate. Mean phase (θ_0_) from 2 independent biological replicates is shown.

**Supplementary Figure 4.**
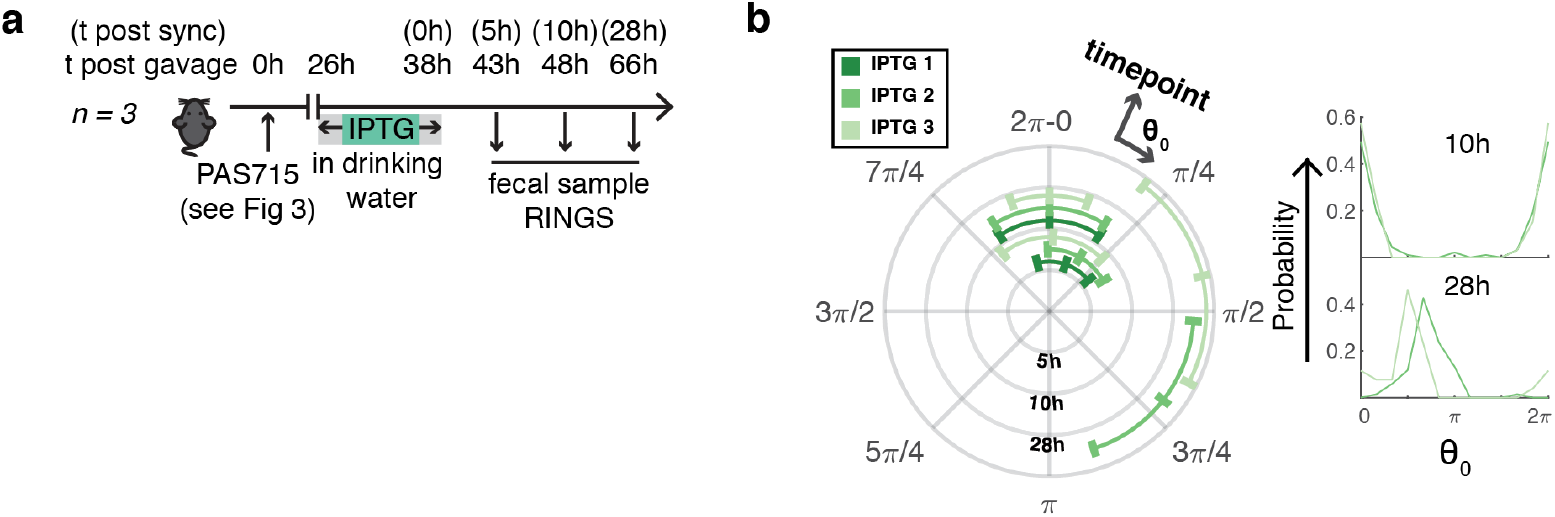
**a.** Following growth analysis in the streptomycin treated gut (Fig 3a,b,d) PAS715 bacteria were re-synchronized by provision of IPTG (lOmM) in 5% sucrose drinking water for ~12 hours overnight. IPTG was then removed and normal drinking water provided, with fecal samples plated and analyzed by RINGS over the following ~24 hours to follow repressilator phase progression. **b.** RINGS analysis showed unchanged phase of the population 5- and 10-hours after removal of IPTG, but clear phase progression 28-hours after IPTG removal. Circular graph (left) shows circular mean ± angular deviation for colonies from each mouse. Growth corresponds to a clockwise phase shift and each ring corresponds to a discrete timepoint taken. Linear histograms (right) compare bacterial phase (θ_0_) distributions at 10- and 28-hours after IPTG removal. Data represent normalized counts from π/6-width bins of phase values.

**Supplementary Figure 5.**
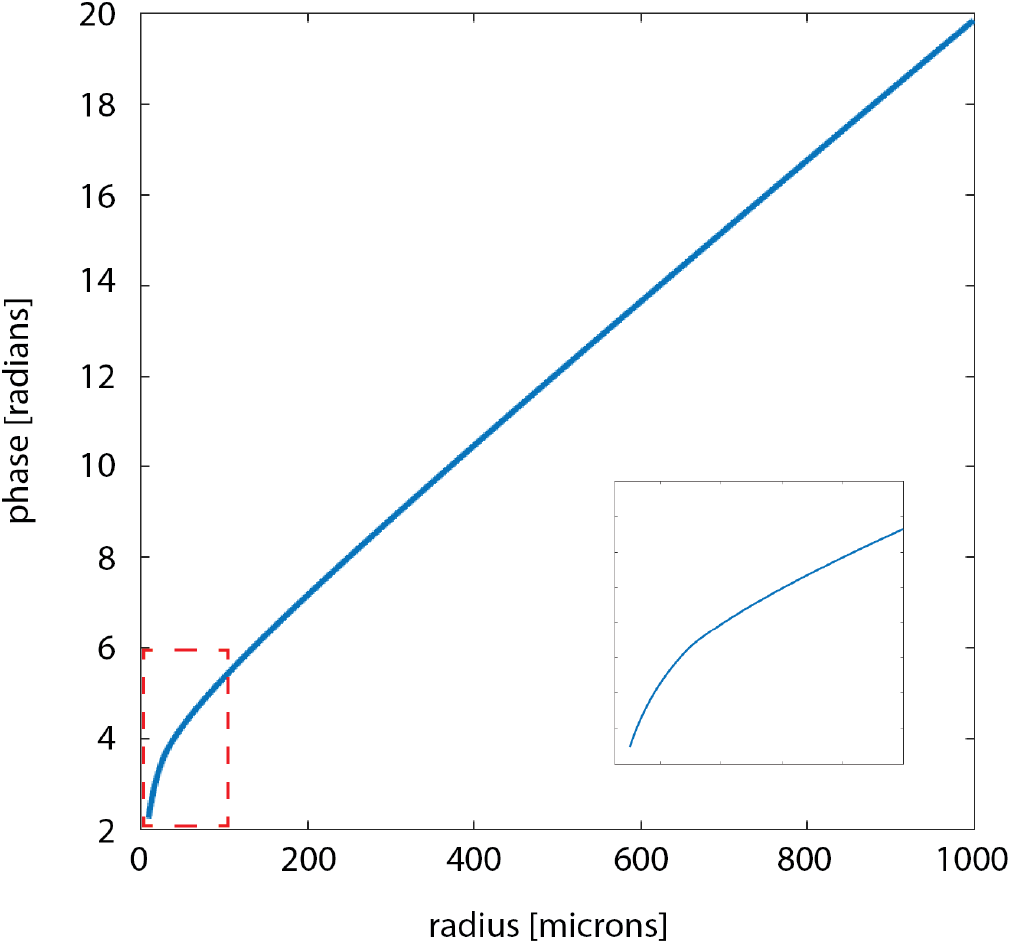
A model for bacterial colony growth. Phase profile of a combined growth model based on an exponential growth phase at low radius and linear growth phase at high radius. Inset shows region marked in red.

**Supplementary Figure 6.**
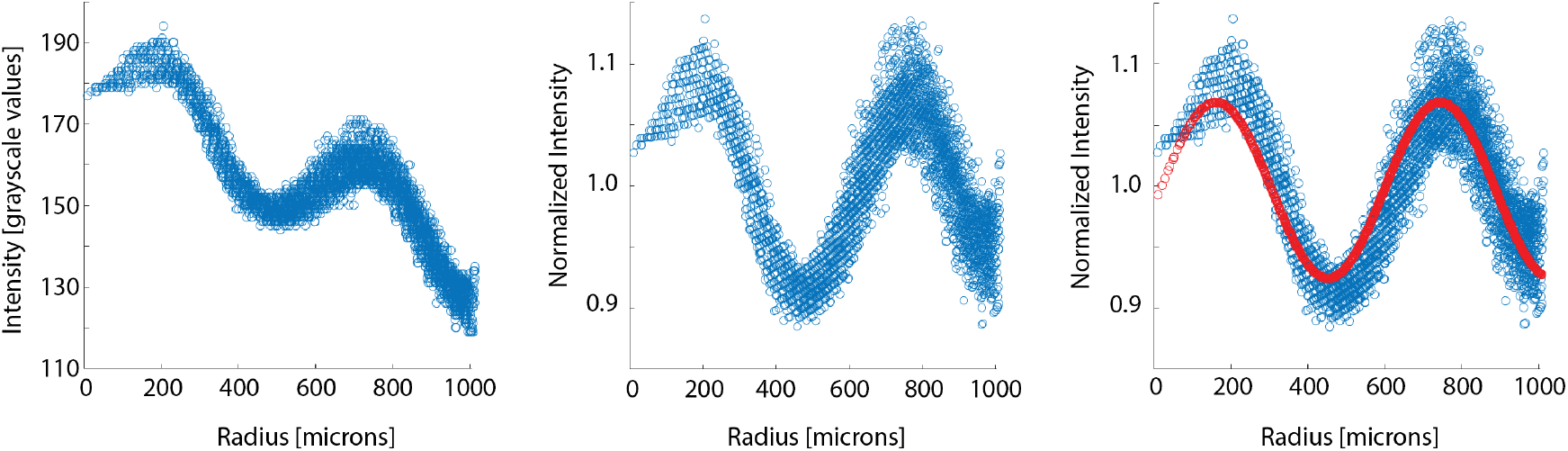
Data normalization and curve fitting by RINGS. **a**. The raw intensity values from the image of a single colony, demonstrating intensity loss at increasing radius. **b**. Intensity values after normalizing by a second order polynomial model. **c**. The model fit (red) overlaid on raw data (blue).

**Supplementary Figure 7.**
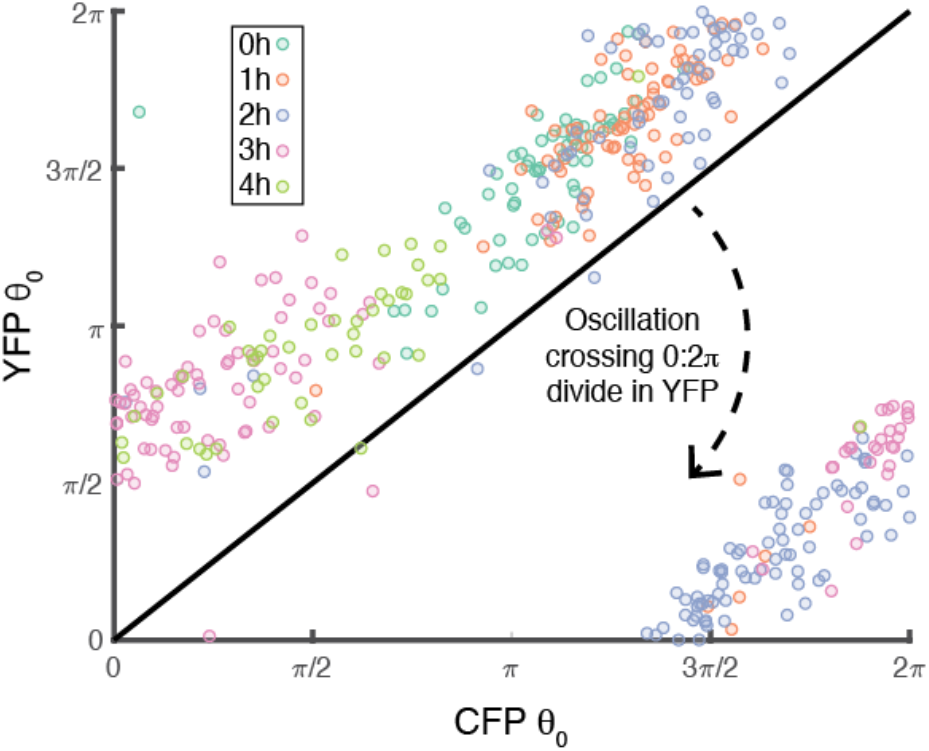
Two-color RINGS analysis. Independent RINGS analysis on YFP and CFP images from a single timecourse of LPT320 bacteria growing in exponential phase on LB medium at 37°C. The constant offset between YFP θ_0_ and CFP θ_0_ is evident in the shift from the 1:1 curve. Also of note is the tendency for correlation between YFP and CFP estimates, even within individual, synchronous timepoints, suggesting that single-channel measurement errors are not the predominant cause of variability within the population after synchronization.

**Supplementary Figure 8.**
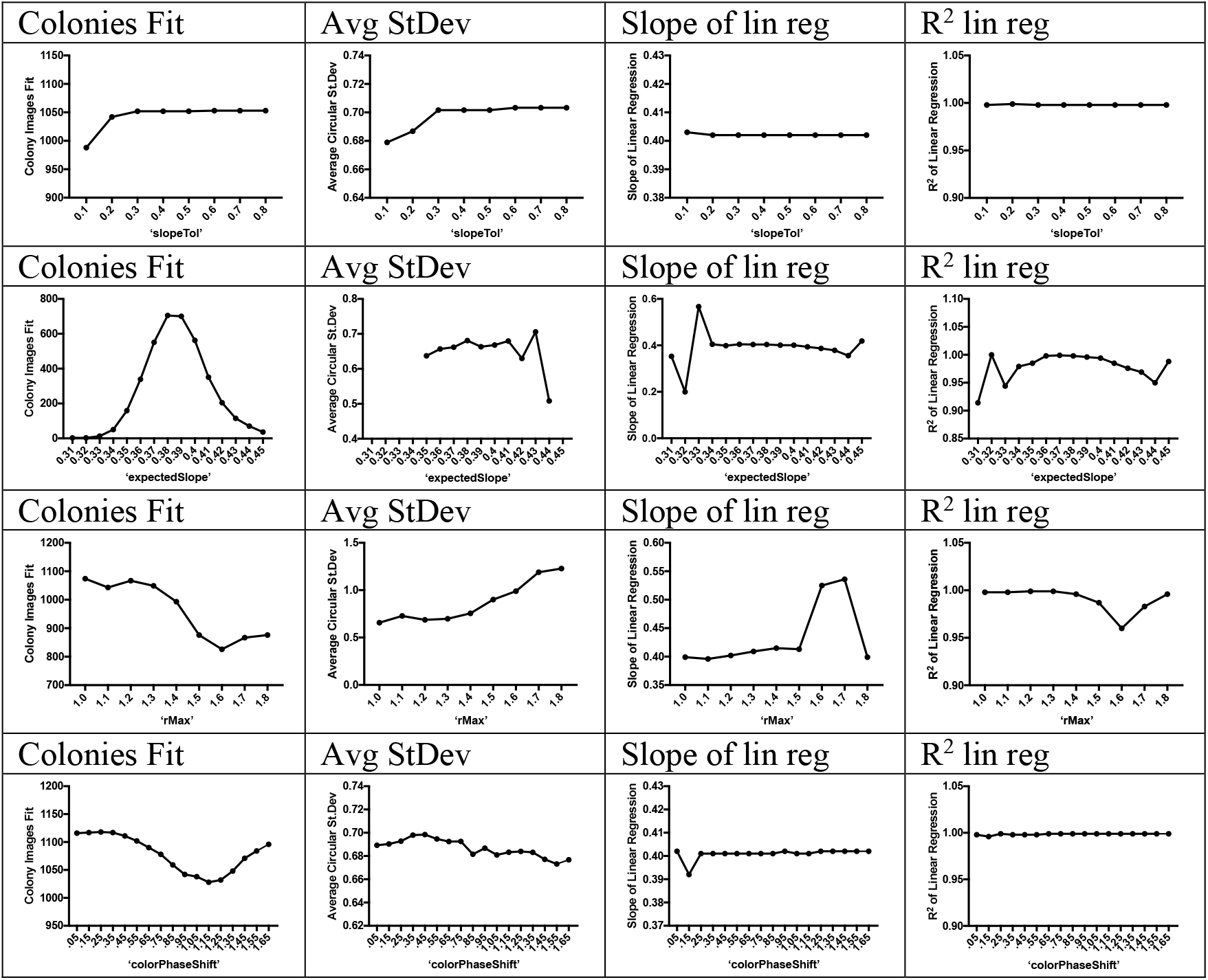
Parameter sweep across key variable parameters for RINGS model fitting. Each parameter was tested on a small timecourse dataset (shown for LPT320) and evaluated based on the total numbers of colonies successfully fit, the average standard deviation across each timepoint, the slope of a linear regression taken through the entire dataset vs generations elapsed (as calculated by CFU counts), and the R^2^ of that dataset fit. Results were similar across a broad set of reasonable parameters. By example, ‘rMax’ variation (3^rd^ line of data) caused little effect to overall fit for all values <1.5. The large effect of values > 1.6, likely corresponding to the point at which the mask exceeded the bounds of colonies within the dataset causing spurious fitting. Ideal parameters were thus calculated from parameter sweeps such as this for each strain.

**Supplementary Figure 9.**
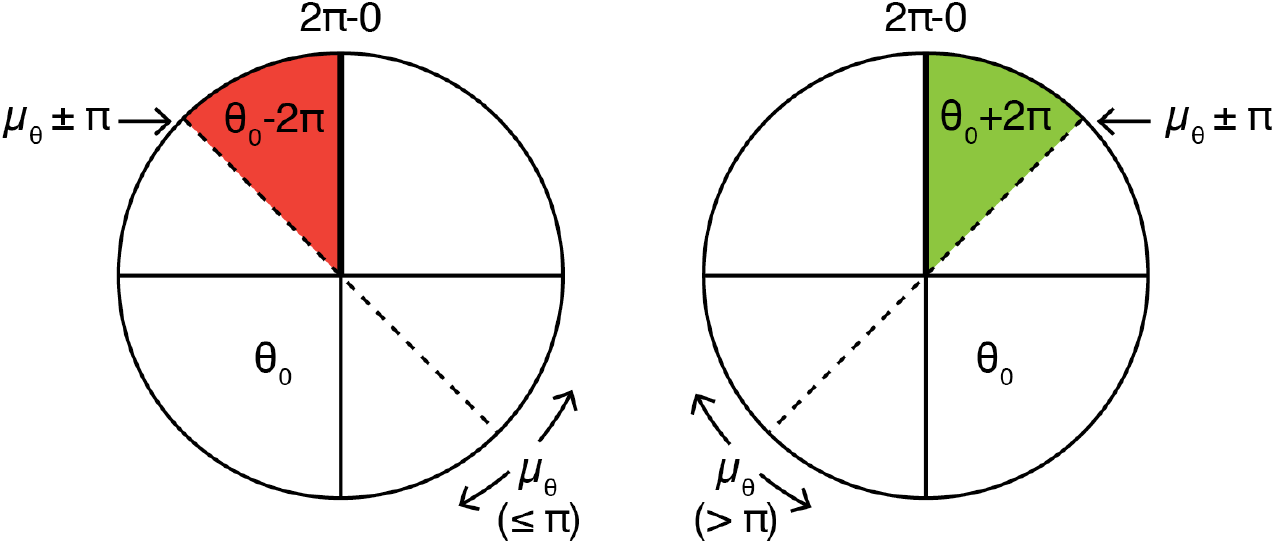
Modulation of phase values for elapsed phase estimation. For elapsed phase calculations, individual phase values (θ_0_) were normalized to the mean (μ) θ_0_ at the starting timepoint (t_sync_). Elapsed phase was then assumed by addition/subtraction of multiples of 2π based on the assumption that all datapoints remained with ± π of the timepoint mean. Where the modulo 2π mean θ_0_ of a timepoint was < π, all values 0 to μ+π were assumed to be one phase revolution (i.e. 2π radians) ahead of values μ-π to 0. Where the modulo 2π mean θ_0_ of a timepoint was >π, values 0 to μ-π were assumed to be one phase revolution (i.e. 2π radians) ahead of values μ-π to 0.

**Supplementary Figure 10:**
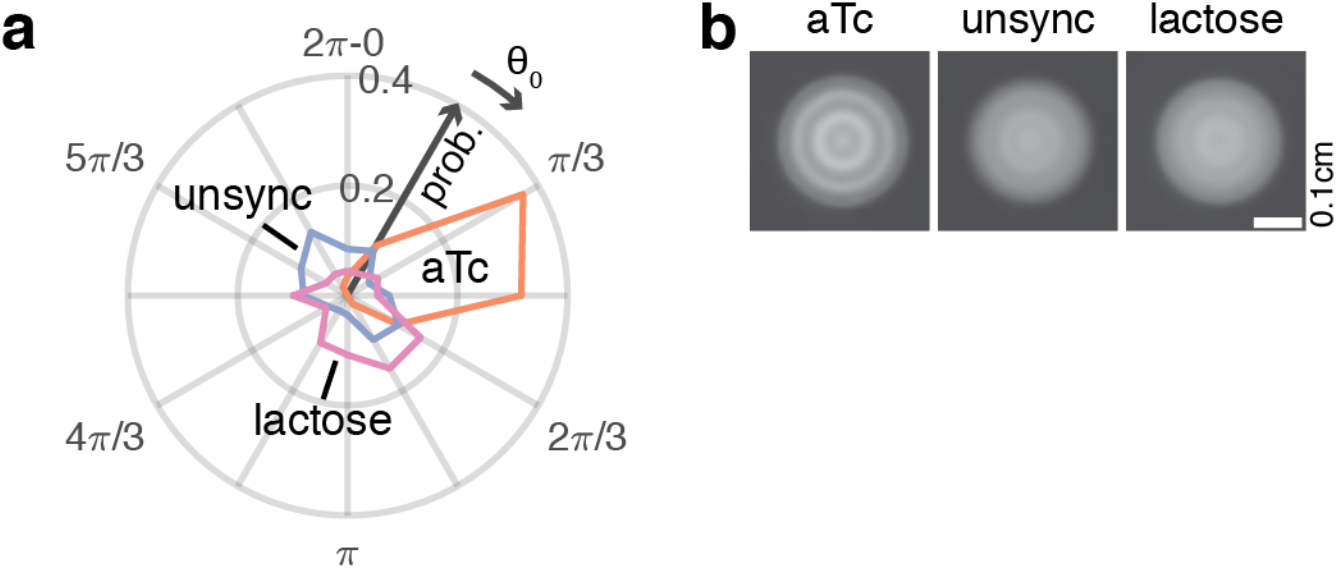
The repressilator is unaffected by growth on lactose. **a.** Growth of LPT239 bacteria, carrying the repressilator, in the presence of 100nm aTc or 0.8% lactose, followed by RINGS analysis, demonstrates the lack of repressilator synchronization caused by lactose. Graph shows polar histogram representing (radius) normalized counts of colonies within a (angle) given θ_0_ range, with π/6-width bins. **b.** Average projections of aligned mVenus fluorescence colony images for each population also demonstrate the lack of synchrony compared to aTc synchronized, and unsynchronized controls. Scale = 0.1cm.

**Supplementary Table 1:** Plasmids used in this study

| Plasmid name | Source | Details |
| --- | --- | --- |
| pLPT234 | This study | pSC101, Amp, repressilator 2.0 plasmid<br>PL<sub>tetO1</sub>-*cl*, PR-*lacI*,<br>PL<sub>lacO1</sub>-*tetR*, PL<sub>tetO1</sub>-*venus*,<br>PL<sub>lacO1</sub>-*cfp*, PR-*mKate2* |
| pLPT41 | <sup>8</sup> | ColE1, Kan, P<sub>tetO1</sub> sponge<br>plasmid |
| pLPT149 | <sup>8</sup> | ColE1, Kan, P<sub>tetO1</sub>+P<sub>R</sub><br>sponge plasmid |
| pLPT145 | <sup>8</sup> | ColE1, Kan,<br>P<sub>tetO1</sub>+P<sub>R</sub>+P<sub>lacO1</sub> sponge<br>plasmid |

**Supplementary Table 2:** Strains used in this study.

| Strain | Source | Details |
| --- | --- | --- |
| LPT239 | This study | *E. coli* MC4100 + pLPT234 + pLPT145 |
| LPT320 | This study | *E. coli* MC4100 + pLPT234 + pLPT41 |
| LPT322 | This study | *E. coli* MC4100 + pLPT234 + pLPT149 |
| PAS715 | This study | *E. coli* MG1655 DE(*lacI*) DE(*motA*) *rpsL* K42R + pLPT234 + pLPT145 |
| PAS716 | This study | *S. Typhimurium* LT2 + pLPT234 + pLPT145<br>(*streptomycin resistant through uncharacterized mutational selection). |
| PAS717 | This study | *E. coli* Nissle 1917 DE(*lacI*) DE(*motA*) *rpsL* K42R + pLPT234 + pLPT145 |
| PAS718 | This study | *E. coli* MG1655 DE(*lacI*) DE(*motA*) *rpsL* K42R<br>attTn7::pRNA1-mKate2 + pLPT234 + pLPT145 |

